# 4 SPECIES OF BACTERIA DETERMINISTICALLY FORM A STABLE BIOFILM IN A MILLIFLUIDIC CHANNEL: ASSEMBLY PRINCIPLES

**DOI:** 10.1101/2021.04.16.440159

**Authors:** A. Monmeyran, W. Benyoussef, P. Thomen, N. Dahmane, A. Baliarda, M. Jules, S. Aymerich, N. Henry

## Abstract

Multispecies microbial adherent communities are widespread in nature and organisms but the principles of their assembly and development remain unclear. Yet, the demand to understand and predict the responses of such living communities to environmental changes is increasing, calling for new approaches. Here, we test the possibility to establish a simplified but relevant model of multispecies biofilm in a laboratory setup enabling *in situ* real-time monitoring of the community development and control of the environmental parameters in order to decipher the mechanisms underlying the formation of the community. Using video-microscopy and species combinatorial approach, we assess the global and individual species spatiotemporal development in millifluidic channels under constant flow of nutrients. Based on quantitative measurements of expansion kinetics, local dynamics and spatial distribution, we demonstrate that the four chosen species (*Bacillus thuringiensis, Pseudomonas fluorescens, Kocuria varians* and *Rhodocyclus* sp.) form a dynamical community that deterministically reaches its equilibrium after about 30 hours of growth. We evidence the emergence of complexity in this simplified community as reported by spatial heterogeneity rise and non-monotonic developmental kinetics. We find interspecies interactions consisting in competition for resources — in particular oxygen — and both direct and indirect physical interactions but no positive feedback. Thereby, we introduce a model of multispecies adherent community where effective couplings result from individual species quest for fitness optimization in a moving and heterogenous environment. This control and the understanding of this simplified experimental model shall open new avenues to apprehend adherent bacterial communities behavior in a context of rapid global change.

## Introduction

Microbial communities are essential for global geochemical cycles and ecological equilibria e.g. (1). In nature, these communities typically comprise hundreds of distinct taxa with an enormous potential of direct and indirect interactions (2). These ecosystems arouse increasing scientific interest in the perspective of predicting and mitigating their response to global environmental change (3). High-throughput sequencing and meta-omics approaches coupled with computational models provide invaluable insights in the microbial taxonomic and functional structure of the complex communities harvested from their natural habitats — soils or oceans for instance (4–6). They make the case that multiple examples of ecological interactions such as syntrophy (7), catabolic parasitism (8) or competition for nutrient resources (9) occur in these microbial associations. Yet, the mechanistic understanding of the dynamics driving the community formation remains limited from these approaches due to the high complexity of a natural system. In addition, the inference of predictive networks in natural bacterial communities faces serious hurdles. Environment spatial heterogeneity is essentially overlooked while it is expected to play a significant role in the community structure and functioning (10). The appropriate time scale is also hard to capture. In particular, whether the community experiences a transient phase or a stationary state at the considered time points is usually unknown. Eventually, the environmental factors are intrinsically impossible to control.

Conversely, experimental models of microbial communities, highly simplified and limited in size, offer the advantage of enabling micro-organism population manipulation, traits and environmental parameters control (3, 11) as well as the completion of experimental replicates. As already argued (11, 12), these laboratory microcosms do not intend to reproduce natural communities but rather to uncover general principles operating in field communities as exemplified in the pioneer work of G.F. Gause (13) who, based on yeasts or paramecia co-cultures, described and conceptualized the interspecies interaction mechanisms that led to his theory of competitive exclusion.

Bacterial communities principally live attached to surfaces in nature, forming cell assemblages embedded into an extracellular polymer matrix that ensures their confinement and supports their structure, with major consequences for their development and properties (14–18). These adherent communities, so-called biofilms, are heterogeneous, highly concentrated and the site of multiple physico-chemical gradients (19). The resulting spatial organization (10, 20–24), is completely lost when samples are harvested from their initial development site to be analyzed in the laboratory. Recent advances in microscopy imaging (25, 26) and microfluidics (27–29) now facilitate biofilm *in situ* explorations and spatial features investigations. Recently, an increasing interest for multispecies systems has emerged. Experimental approaches dedicated to multispecies biofilms also begin to report a whole range of social interactions and spatial structures in specific models (30–36). However, due to the difficulty to implement accurate real-time imaging in composite systems, in most of these studies, biofilms are imaged at given time points using FISH and adaptations of the technique on fixed samples (22, 33, 37–44), which misses or blurs kinetic information. On the other hand, individual-based or continuum modeling approaches help to formalize mechanisms potentially involved in biofilm structure (45–49). Yet, the complexity of an adherent community development engages many processes combining cell biological traits and environmental physicochemical properties impossible to fully integrate in the model.

Hence, we reason that a kinetic analysis of the formation of a multispecies biofilm should provide more insights into the mechanistic bases underpinning community establishment and dynamics. For this work, we build an experimental model composed of four species of bacteria derived from a natural biofilm which accidentally developed in the industrial context of a pasteurization line (50). Our assembly which included *Bacillus thuringiensis (Bt), Pseudomonas fluorescens (Pf), Kocuria varians (Kv)* and *Rhodocyclus* sp. *(Rh)* aimed at a trade-off between simplification, preservation of a sufficient diversity and cultivability. The four species are *a priori* only related by their ability to co-exist spontaneously, which opens a large spectrum of mechanisms potentially behind the formation of the community. Our setup places the four species in a millifluidic channel under flow (51), realistic to a whole range of living or abiotic environments such as veins or streams (2). Made of PDMS and glass, the device provides regulated hydrodynamics and medium supply. In the presence of a growing biofilm, physical and chemical gradients are generated, offering a controlled habitat, complex enough to create environmental diversity. The spatiotemporal development of the composite adherent community is monitored using optical video-microscopy. We also implement a species combinatorial approach which implies building all the different combinations of the 4 species to decipher the full community specific traits.

Thus, we demonstrate that the four chosen species form a dynamic adherent community that deterministically reaches its equilibrium after about 30 hours of growth. We provide evidences for the emergence of spatial heterogeneity and non-monotonic developmental kinetics attesting for the rise of complexity in this simplified model based on competitive and physical interspecies interactions. We discuss the community mode of organization of this experimental model in the perspective of further investigations on the response of this system to perturbations.

## Materials and Methods

### Bacterial strains and culture conditions

#### Bacillus thuringiensis

(*Bt*) is a 407 Cry^−^ strain (52). The fluorescent variants — *Bt*-FAST and *Bt*-GFP — were genetically engineered to express either the protein FAST (53) or the protein GFP under the control of the constitutive promoter P*sar. Pseudomonas fluorescens* (*Pf*) was an mCherr*y*-expressing strain (WCS365 containing pMP7605)(54), a gift from E.L. Lagendijk from Leiden University (The Netherlands). *Kocuria varians* (CCL56) and *Rhodocyclus sp*. (CCL5) were isolated from a biofilm formed on a gasket in a milk pasteurization line (50). The strains were routinely cultivated at 30°C on M1 medium (see Supplementary information).

### Millifludidic device

We micro-fabricated millifluidic channels 30 mm in length, 1 mm in width and height. A polydimethylsiloxane (PDMS) mixture (RTV615A+B from Momentive) was poured at ambient temperature in a polyvinyl chloride home-micromachined mold and left to cure at least 3 hours in an oven set at 65°C. Then, the recovered templates were drilled for further plugging of adapted connectors and tubings. PDMS templates and glass coverslips were then cleaned using an oxygen plasma cleaner (Harrick) and immediately bound together to seal the channels. The last step consisted in adapting connections: we used stainless steel connectors (0.013” ID and 0.025” OD) and microbore Tygon tubing (0.020” ID and 0.06” OD) supplied by Phymep (France). The thin metallic connectors accommodate a bottleneck on the flow circuit which prevented upstream colonization. The medium was pushed into the channels at a controlled rate using syringe pumps for the 36 to 40 hours of the experiment. Up to 12 channels could be run and monitored in parallel. The whole experiment was thermostated at 30°C.

### Biofilm formation

#### Initiation

The same number of cells of each species — approx. 1.2×10^5^ cells — of exponentially growing cultures were mixed and immediately injected in the channels directly in the PDMS channels using a syringe with 22G needle before connecting the tubings. Then, the cells were allowed to settle down for 1h30 before starting medium flow. Nutrient flow triggering was taken at time t=0. For biofilm growth, we used MB medium, an adaptation of M1 medium. Additional details are given in Supplementary information.

#### Attached community development

The flow rate of 1 mL/h imposed a mean advection characteristic time *τ*_*a*_ of 2 min while in the stokes-Einstein approximation the characteristic sedimentation time *τ*_*s*_ is in the order of the hour and the diffusion characteristic time *τ*_*d*_ of the order of the hour for motile bacteria to tens of hours for non-motile cells (see details in Supplementary Information). This implies that, on average, bacteria suspended in the channel will be continuously quickly washed-out by the flow, so that essentially attached cells reside and divide in the channel together with a minority of freshly detached cells. Microscope time-lapse imaging of the channel bottom surface was started a few minutes before the triggering of the flow at time t=0.

### Microscope imaging

#### Microscope

We used an inverted NIKON TE300 microscope equipped with motorized x, y, z displacements and shutters. Images were collected using a 20 × S plan Fluor objective, NA 0.45 WD 8.2-6.9. Bright field images were collected in direct illumination (no phase). Fluorescence acquisitions were performed using either the green channel filters for GFP and FAST:HBR-2,5-DM (Ex. 482/35, DM 506 Em. FF01-536/40) or the red one for m-Cherry (Ex 562/40nm DM 593 Em. 641/75). Excitation was performed using a LEDs box (CoolLed pE 4000).

#### Epifluorescence

We collected the fluorescence signals focusing the image on the bottom surface. Owing to the small numerical aperture (0.45) of the objective (20x) and the 1 mm-height of the channel, we collected the signal from the whole channel height (Fig. S1)

#### Confocal microscopy

For specific purpose, we also collected confocal images using a spinning disk Crest X light V2 module (Gataca, France distribution) exhibiting an axial resolution of 5.8 *µ*m.

### Image acquisition

We used a Hamamatsu ORCA-R2 EMCCD camera for time-lapse acquisitions of 1344×1024 pixels images with 12 bits grey level depth (4096 grey levels) and captured an *xy* field of view of 330 *µ*m × 430 *µ*m. Bright field and fluorescence images were usually collected for 36 hours at the frequency of 6 frames per hour. Excitation LEDs were set at a 50% power level and exposure times were 50 ms or 500 ms for the green emissions — GFP and FAST respectively — and 800 ms for the red emissions.

### Image analysis

#### Image intensities

Timelapse images were analysed to derive the kinetics of biomass accumulation in the channel based on microscopic optical density (*µ*OD) measured from transmitted light images according to *µ*OD = ln (*I*_*0*_/*I*), where *I*_*0*_ is the intensity recorded on a channel filled with medium only and *I* the intensity recorded on a channel containing a growing biofilm (51). Image intensity per pixel averaged on the whole image or on defined regions of interest (ROIs) was collected using the NIKON proprietary software NIS. The data sheets edited by NIS were next exported to Matlab for further analysis of the biofilm development kinetics and growth parameters determination. *Bt* and *Pf* expansion kinetics was measured from timelapse images fluorescence intensities in their respective optical channel. Background was subtracted using the contribution to the fluorescence intensity of a channel of medium in the absence of bacteria. All curves are averaged over at least three independent replicates.

#### Kv delineation

*Kv* cells and clusters in the community were delineated using transmitted light image thresholding and morphological descriptor filtering from using NIS smart thresholding tool (see Fig. S2 for more details).

#### Pearson’s correlation coefficient

Fluorescent species dynamics was evaluated calculating *r*_*c*_, the Pearson’s correlation coefficient from consecutive frames of the growing-biofilm movies using ImageJ 1.52k correlator (55). This coefficient is equal to 1 for perfectly identical images and close to 0 for completely distinct ones (Fig. S3).

## Results

### The 4 species assemble according to a robust temporal sequence and form a community ultimately reaching dynamic equilibrium

#### Community formation

To assess the global development of the attached community under constant flow of growth medium in the millifluidic channel (Fig. 1A), we acquired time-lapse images (Fig. 1B) and measured the temporal variations of the biomass as reported by the microscopic optical density (Fig. 1C). The kinetics shows a biomass increase that levels off after approx. 25 hours. In addition, several inflections, robust across positions in the same channel, channels and biological replicates, attested for the succession of 2 characteristic temporal phases with climaxes sharply delineated by the derivative curve which measures biofilm expansion rate (Fig. 1D). A first biofilm initiation phase spread over the first 8 hours and included an accelerated growth phase (increasing expansion rate) followed by a damping phase (decreasing expansion rate) with a climax at t = 5 hours. Next, a second expansion phase arose exhibiting a climax around t = 16 hours followed by a kinetic steady-state. Remarkably, the stabilization of the average level of biomass coincided with a marked increase of the noise in biomass (Fig. 1C). Movies built from the image stacks clearly show that the noise came from the frequent passage of biofilm flocs detached upstream of the observed position (Movie S1). Locally, detachment events can be observed creating gaps refilled in a few tens of minutes (Movie S2). Thus, the kinetic steady-state signal likely results from the balance between biofilm growth and biofilm detachment, consistently with a dynamic equilibrium.

**Figure 1:**
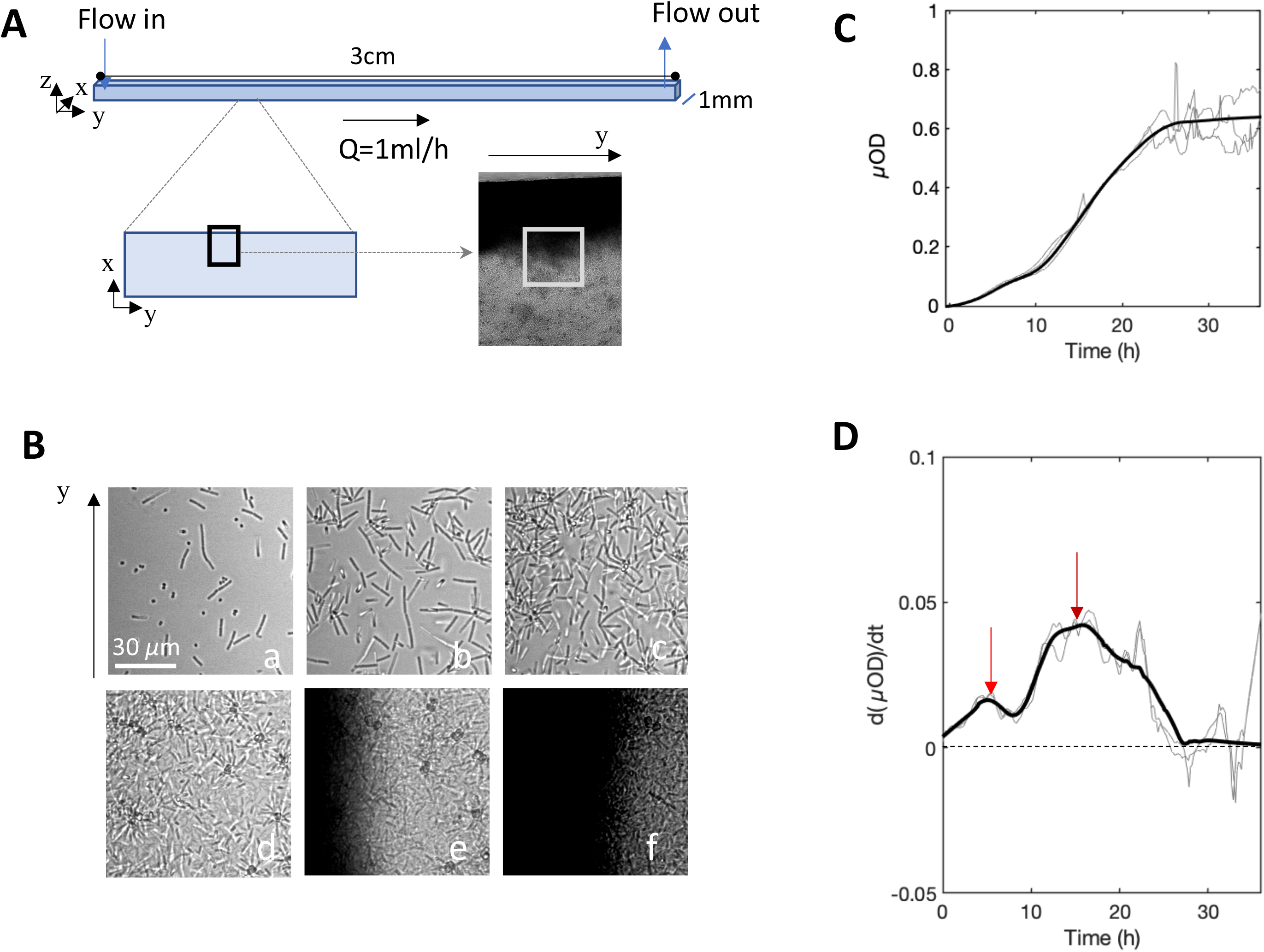
4S biofilm growth under constant flow in the millifluidic channel. **(A)** A square channel of 1mm side and 30 mm long continuously fed with growth medium at 1 mL/h is settled on the stage of a microscope thermostated at 30°C. Images of 0.15 *µ*m^2^ size, covering near half of the width of the channel from the edge to the middle (black rectangle) are taken every 10 mins. (**B**) Zooming in on center of these images (grey square in panel A) at time t = 10 min (a); 2h30 (b); 5h (c); 18h (d); 30h (e); 35h (f). (**C**) *µ*OD — calculated from transmitted light images, *i*.*e*. ln(*I*_0_/*I*), and proportional to biomass — as a function of time. The intensity *I* is averaged over all the pixels of the 0.15 *µ*m^2^ image (black frame in panel A). (**D**) Derivative of *µ*OD signal with respect to time. The red arrows show maxima of the expansion rate. Light curves are from three independent biological replicates, the bold curve is the smoothed average.

The images (Fig. 1B) revealed the evolution of the spatial distribution of the biofilm in the channel. Initially, the colonization occurred and progressed uniformly on the bottom surface. Then, in the second expansion phase, an accumulation at the channel edges emerged while the central zone remained less densely populated.

#### Steady-state assessment

In order to assess the stability of the community steady-state, we evaluated the ability of the community to recover after a perturbation. To this purpose, at time t = 38 hours, we applied a major physical disturbance by injecting a 200 µl air bubble which detached approx. half of the attached biomass, indeed *µ*OD was reduced by a factor of 2 bringing back the biofilm to the level it reached 20 hours earlier. The results in Fig. 2 show that the community biomass returned to its previous steady-state (before perturbation) in approx. 20 hours, suggesting the formation of a stable community (56, 57). Yet, the highest signal-to-noise ratio of the recovery part of the curve which corresponds to small floc detachment might reveal a slightly higher physical fragility of the material re-formed after the physical perturbation.

**Figure 2:**
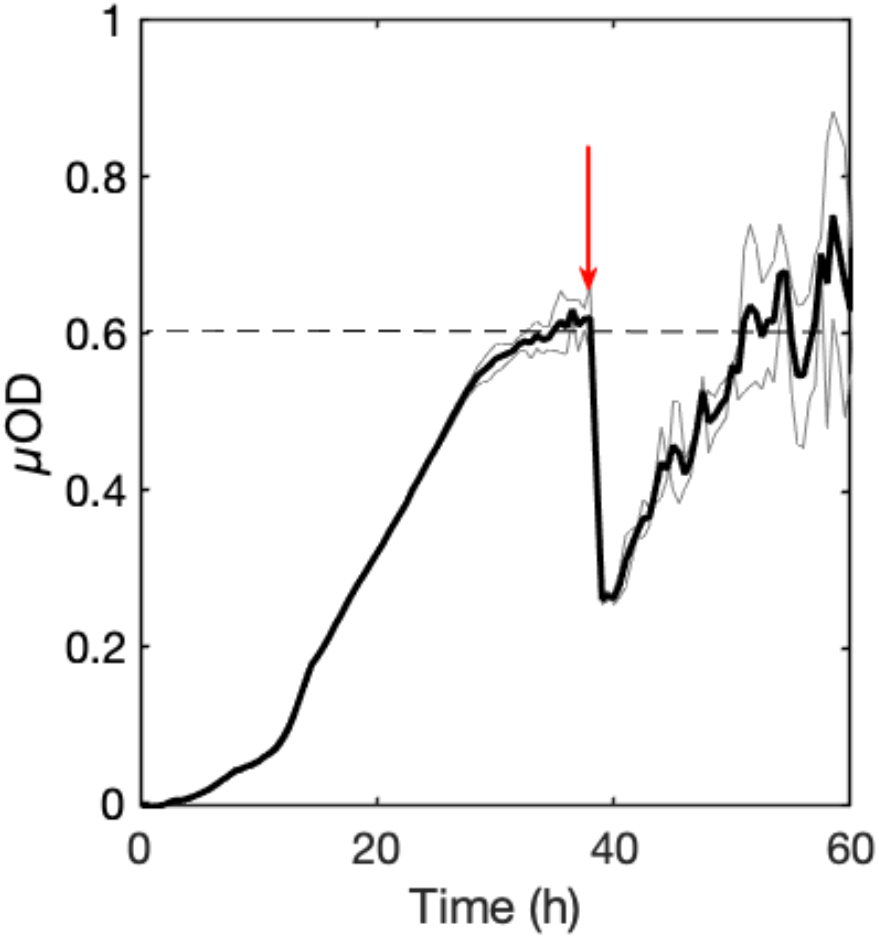
4S biofilm recovery after perturbation. Evolution of the biomass (as reported by *µ*OD) of the 4S biofilm grown in the 1 mm-side square channel at 1 mL/h for 38 hours at 30°C, then partially wrecked by the injection of a 200 µL-bubble of air (red arrow). The bold curve shows the average of 3 distinct positions in the channel (light grey curves).

### Species-specific signal delineates individual kinetics and suggests interspecies coupling

To decipher the features of the 4-species (4S) biofilm development, we scrutinized the discrete contribution of each species to the community formation. To this purpose, we exploited *Bt* and *Pf* fluorescence signals (Fig. 3 A and B), *Kv* high contrast in transmitted-light mode (Fig. 3C) and we counted *Rh* cells at final time points after removing the biofilm from the channel (Fig. 3D). *Bt* and *Pf* fluorescence kinetics exhibited multiphasic curves composed of an initial oscillation followed by an intensive growth phase tending to a plateau as already detected on the global transmitted light signal. The derivatization of the kinetics featured the succession of the distinct phases and accentuated the main climaxes (Fig. 3 A and B) enabling extraction of three characteristic times for each species: *t*_1_, the maximum of the expansion rate (beginning of the dampening); *t*_2_, the end of the initial oscillation taken as the point where the derivative becomes positive again and *t*_3_, the maximum of the recovery corresponding to the inflexion preceding the plateau (table 1). Although *Bt* and *Pf* development followed similar kinetics, we observed a few hours shift in the characteristic times related to each species — intriguingly *Pf* exhibited delayed t_1_ and t_2_ but earlier plateau compared with *Bt*. The correspondence of the *Bt* and *Pf* kinetics suggested the existence of a coupling although climaxes were not fully synchronized.

**Figure 3:**
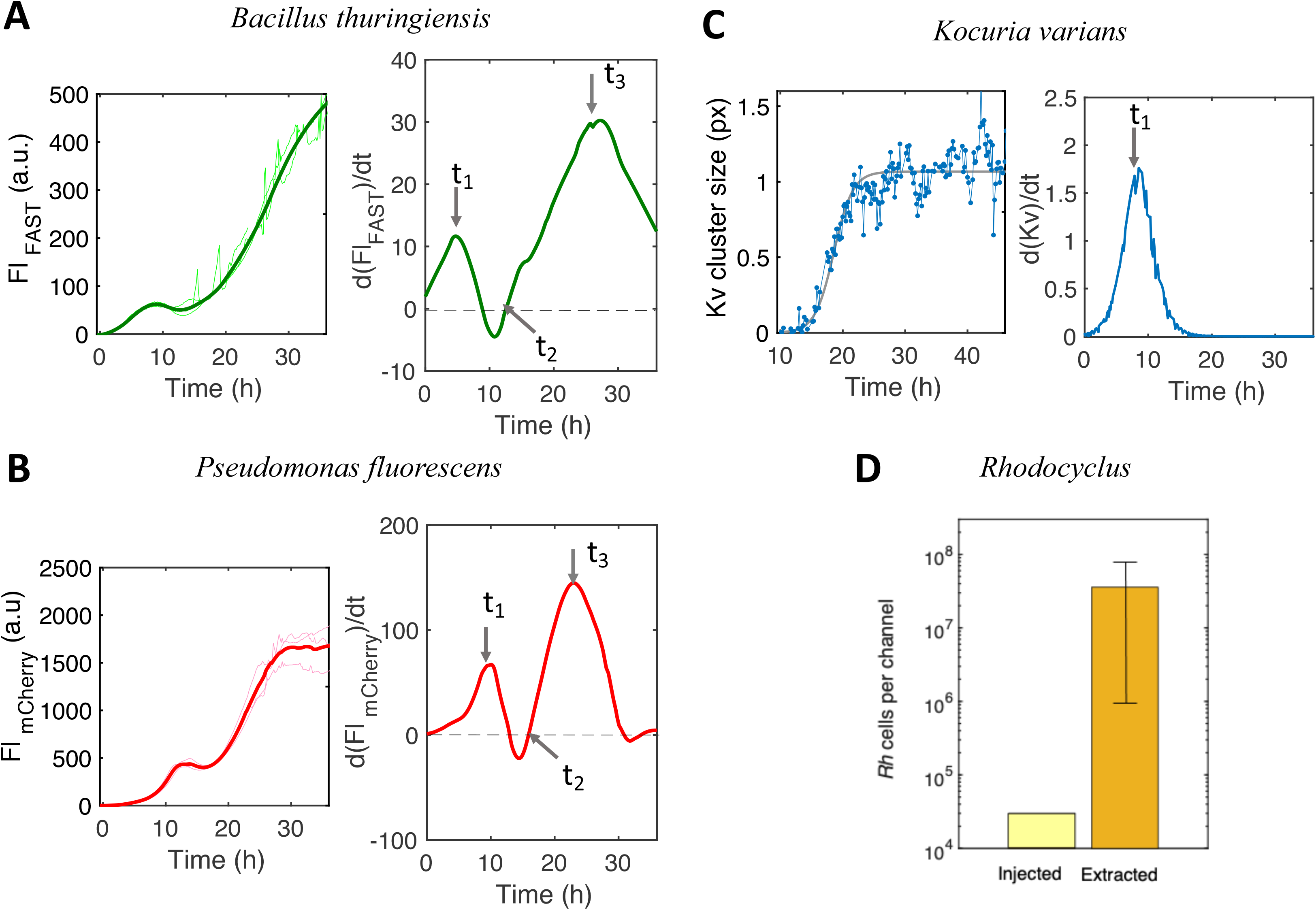
Individual species developments in the 4S adherent community. (**A, B**) Kinetics of the fluorescence intensity, Fl, from *Bt*-FAST (panel A, left side) and *Pf*-mCherry (panel B, left side) in the 4S biofilm with the corresponding Fl curve derivative (panels A and B, right side). Fl is the intensity per pixel averaged over the whole image. The arrows on the derivative curves mark the characteristic times t_1_, t_2_, t_3_ reported in table 1. (**C**) *Kv* area detected on transmitted light images as a function of time (left panel, blue line) and corresponding logistic adjustment (grey line) together with its derivative (right panel) (details in SI, Fig S2). In **A, B** and **C**, signals are collected from 0.15 *µ*m^2^ images located as in Fig. 1A (black frame). Light curves are from 3 independent experiments, the bold curve is the smoothed average. (**D**) Bar graph of *Rh* cells injected in the channel (light yellow bar) and recovered after 36 hours of growth (dark yellow bar). Experimental conditions as in Fig. 1.

**Table I:** Characteristic times of individual species expansion in the 4S biofilm in the whole channel.

| **Species** | **t1<sup>a\*</sup>** | **t2<sup>a\*\*</sup>** | **t3<sup>a\*\*\*</sup>** |
| --- | --- | --- | --- |
| ***Bt*** | **4.9±0.2** | **12.58** | **27.1±0.3** |
| ***Pf*** | **9.5±0.5** | **15.9** | **22.9±0.5** |
| ***Kv*** | **8.5** | **NA** | **NA** |

<sup>a</sup> times in hours
\* climax time determined at the first maximum of the fluorescence intensity derivative curve.
\*\* time of the expansion rate variation inversion, fluorescence intensity derivative equal to zero.
\*\*\* climax time at the second maximum.

In parallel experiments, we monitored the fluorescence signal of *Bt*-GFP in the 4S community and compared it to *Bt*-FAST kinetics in order to get an indirect evaluation of O_2_ depletion along the biofilm formation. Indeed, as shown in a previous work (58), the proportionality between FAST and GFP signals is lost when environmental O_2_ level decreases below the threshold enabling GFP final maturation and fluorescence. We observed that the *Bt*-GFP fluorescence curve, initially superimposable with that of *Bt*-FAST, diverged at time t = 5h (Fig. 4), which corresponded to t_1_, the first *Bt* growth climax.

**Figure 4:**
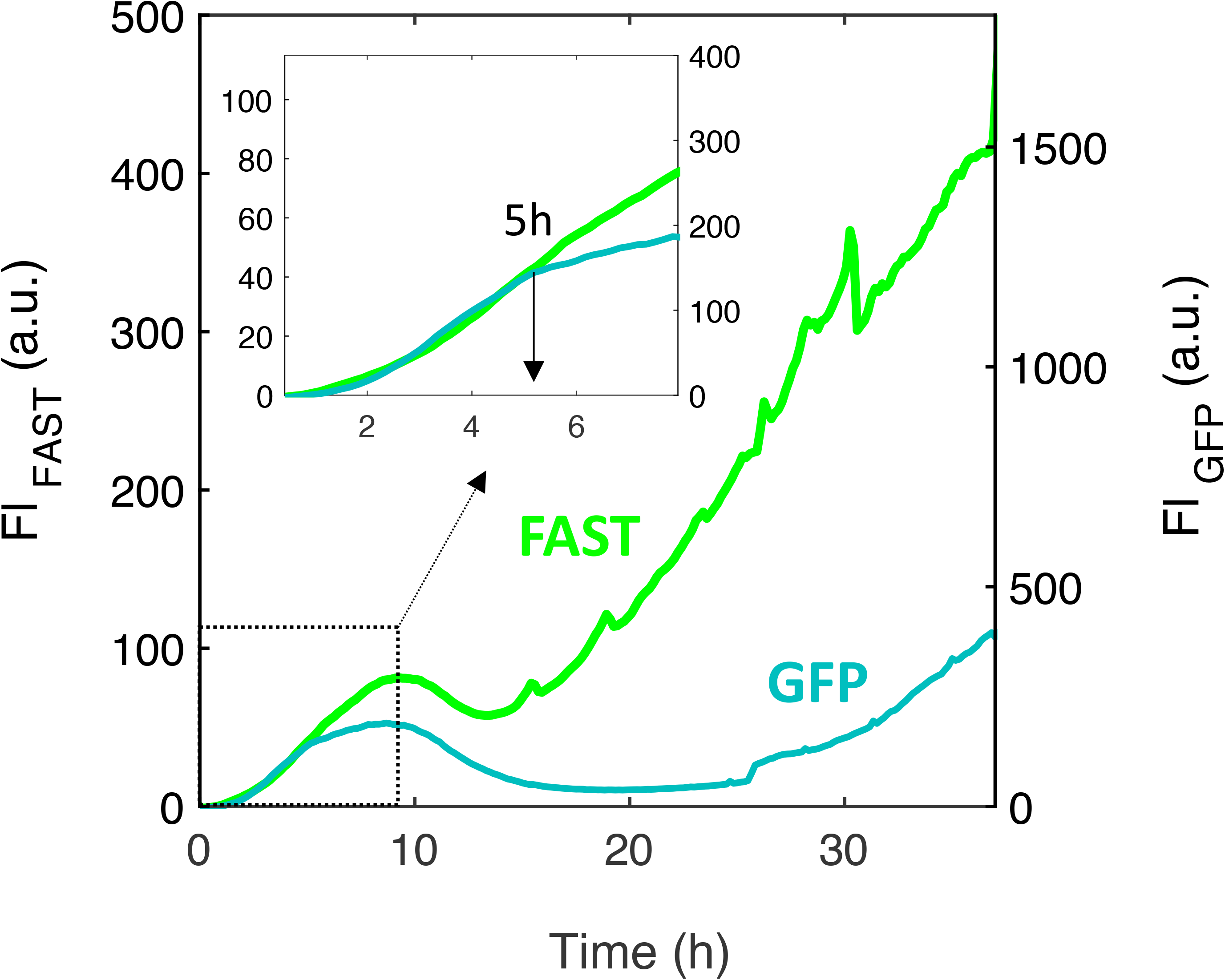
*Bt*-FAST and *Bt*-GFP signals diverge at characteristic time t_1._ Kinetics of the fluorescence intensity, Fl, from *Bt*-FAST and *Bt*-GFP growing in parallel channels in the 4S biofilm. The curves represent the average of at least three independent measurements. The insert shows a zoom in of the first 8 hours of the biofilm formation. Experimental conditions as in Fig. 3A.

In the meantime, the *Kv* population developed from initially attached single cells under the form of small clusters, the size of which was taken as a growth index to plot *Kv* expansion kinetics in the community (Fig. 3C and SI). The obtained curve was formally adjusted to a logistic growth with only one characteristic time t_1_ reporting a single growth phase (table 1). Due to the small cell size, high motility and absence of labeling, the temporal evolution of the *Rh* population could not be accurately determined. Hence, the contribution of *Rh* was evaluated by enumerating the cells after biofilm extraction at the final stage which showed that the *Rh* population successfully thrived in the community, having multiplied by 3 logs along the biofilm development (Fig. 3D). In comparison, *Bt* and *Pf* were multiplied by approx. 3 and 4 logs respectively (Fig. S4).

The results suggest the existence of a coupling between *Bt* and *Pf* populations which both exhibit two successive growth phases, similarly to the global biomass kinetics. By contrast, *Kv* population exhibits an only one-phase expansion. The individual kinetics show that the four species were present all along the biofilm development. An oxygen depletion was deduced in correlation with the first climax of the community.

### The 4S community displays species-specific spatial distribution

To document spatial heterogeneity emergence in the community, we examined the evolution of the species spatial distribution along the community expansion.

We first investigated the species vertical distribution. To this purpose, we run a series of confocal microscopy acquisitions to selectively image bottom and top surface of the channel whereas epifluorescence recordings captured the signal from the whole channel height. We found that *Bt* strictly dwelled on the bottom surface of the channel while *Pf* colonized both bottom and top surfaces (Fig. 5). *Pf* bottom and top surface populations displayed similar kinetic profiles except the first oscillation was delayed by 5 hours on top surface in comparison with the bottom surface. These acquisitions also showed that *Bt* and *Pf* exhibited synchronous kinetics on the bottom surface (see table 1 and table 2). Besides, transmitted light observations showed that *Kv* locations were also limited to bottom surface (Fig. S5). Thus, *Pf* appeared to share the bottom surface with the other species but also to colonize a specific niche on the top surface.

**Figure 5:**
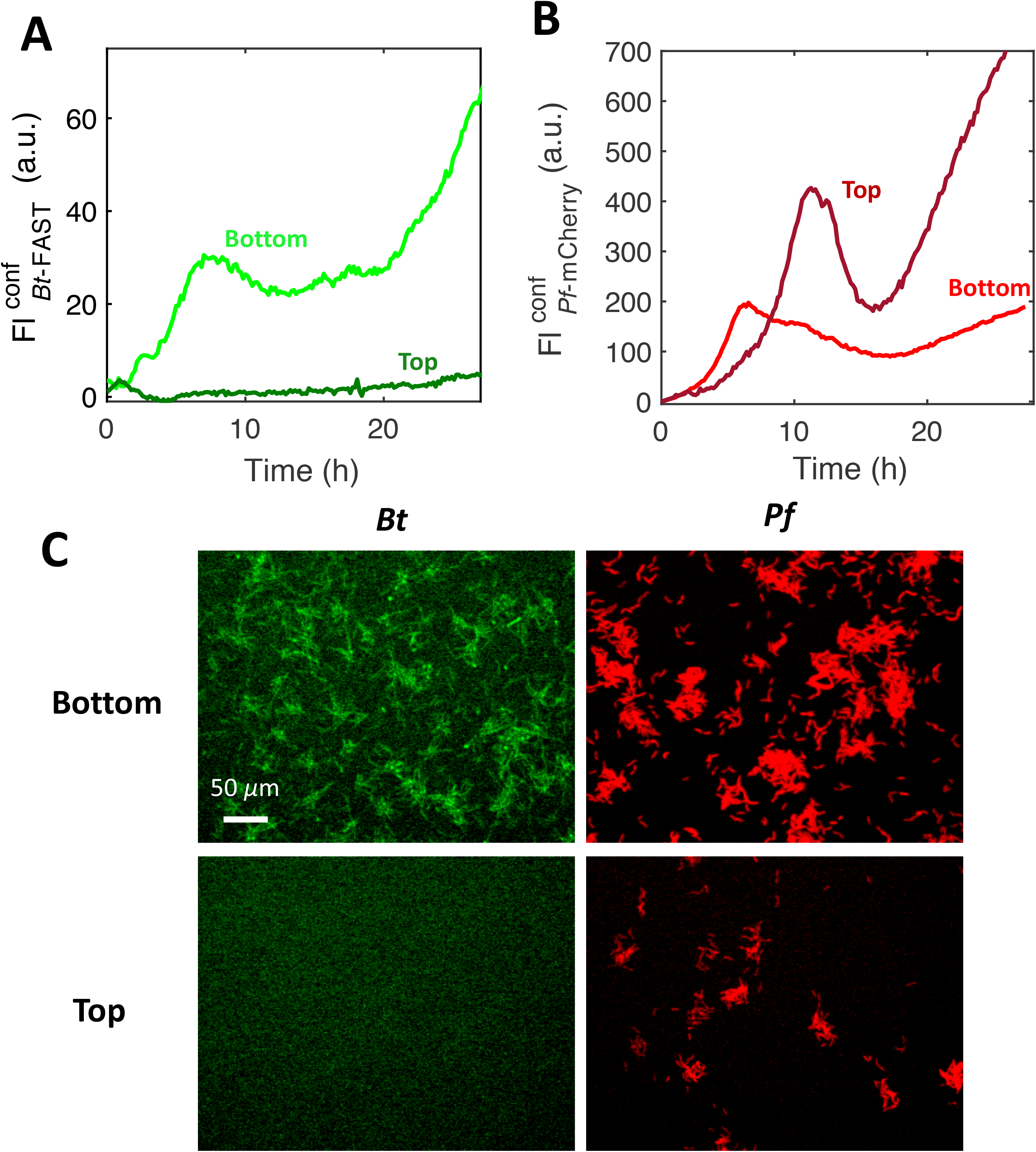
*Bt* and *Pf* dwelling in the channel: bottom versus top surface colonization. Confocal fluorescence intensity of **(A)** *Bt* and **(B)** *Pf* recorded with a focus on the bottom surface (light green **(A)** and light red **(B)**) and on the top surface (dark green **(A)** and dark red **(B)**). Axial resolutions are 5 *µ*m and 5.8 *µ*m for FAST and mCherry signals, respectively. **(C)** Confocal images recorded at time t=3h on the bottom surface (upper panels) and on the top surface (lower panels) for *Bt* (left panels) and *Pf* (right panels). Experimental conditions as in Fig. 1.

**Table II:** Pf bottom versus top surface kinetics.

| Localization | t1<sup>a</sup> | t2<sup>a</sup> | t3<sup>a</sup> |
| --- | --- | --- | --- |
| Pf bottom | 5.04 | 15.92 | 23.08 |
| Pf Top | 11.09 | 16 | NA |

<sup>a</sup> times defined as in table I

Next, we examined the lateral spatial distribution of the individual species, having in mind the spatial distribution shift of the global biomass reported above (Fig. 1B). During the first 5 hours of the colonization, both *Bt* and *Pf* developed a uniform distribution along the x and y axis on the bottom surface before priming a progressive shift towards the edges of the channel (Fig. 6A and B). In order to quantify this shift, we defined two 20*µ*m-wide regions of interest, ROI_Edge_ and ROI_Center_, located at 20*µ*m and 360*µ*m of the channel wall, respectively. Then, we calculated the ratio of their mean fluorescence intensities, FL_Edge/_ FL_Center_ which measured the intensity of the spatial shift. The evolution of this ratio along the formation of the biofilm together with representative kymographs are shown in Fig. 6C. They show a first 5 hours even distribution, then a trend to preferably populate the edge of the channel emerging between 5 and 10 hours and a clear-cut spatial distribution transition toward the channel edge arising shortly after the onset of the second growth phase.

**Figure 6:**
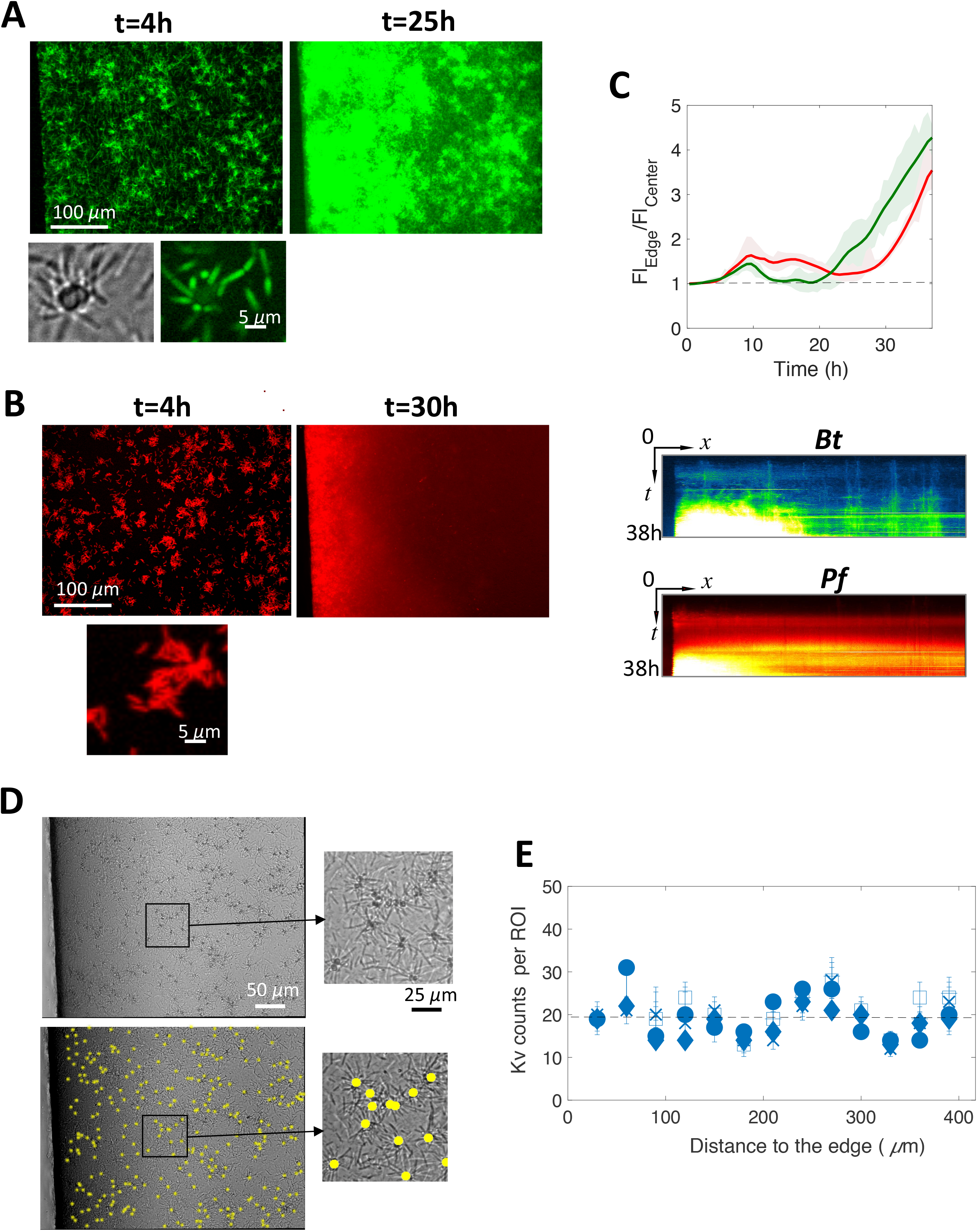
Spatial transition to the channel edge. Fluorescence images of *Bt* (**A**) and *Pf* (**B**) populations in the 4S biofilm, taken in the initial colonization phase (left image) and after the spatial transition (right image). The small bottom images are zoomed-in details showing local heterogeneities of the globally uniform distribution in the initial colonization phase. Graph of the ratio as a function of time of the fluorescence intensity at the channel edge (Fl_edge_) to that at the channel center averaged on ROIs (20*µ*m-wide) located at 20*µ*m and 360*µ*m of the channel wall, respectively; *Bt* in green and *Pf* in red (**C upper panel**). Kymographs of *Bt* (green blue) and *Pf* (Red) spatial distribution (**C bottom panel**). Detection of the Kv cells and clusters in the 4S biofilm using the NIS dark-spot detection tool (details in SI). Images correspond to time t=14h. In the bottom panel, detection results are overlaid on the original image shown in the upper panel (**D**). *Kv* spots are then counted in 30*µ*m-wide ROIs from the edge to the center of the channel for time t= 10 mins (□); 20 mins (•); 7h30 (x); 16h30 (◻); 32h (+); error bars report the error of 10% made on spot counts as evaluated comparing automatic and visual detection.

*Kv* spatial distribution was analyzed collecting on the images the (x,y) coordinates of the *Kv* cells and clusters attached on the channel bottom surface (Fig. 6D and E). We found that the *Kv* discrete locations were evenly distributed over the bottom surface all along the biofilm development — despite short length scale heterogeneities — indicating that *Kv* did not participate to the spatial transition supported by *Bt* and *Pf* consistently with its kinetic profile also clearly distinct from that of *Pf* and *Bt*. The synchronized *Pf* and *Bt* spatial transition reinforces the hypothesis of a coupling of these two populations in the community on the bottom surface. In addition, the temporal correspondence of this spatial transition with the second growth phase indicates that this apparent translocation rather corresponds to a geographically-restricted growth. All the species coexisted on the bottom surface but *Pf* was in addition able to accommodate a private niche on the top surface.

### Local dynamics analysis reveals both static and highly mobile species

To better characterize the initiation phase of the biofilm, we examined the four species local dynamics within the first 6 to 7 hours of the biofilm installation, when the colonized fraction of the surface is small enough to enable individual objects delineation — a period of time corresponding to the first growth phase.

The dynamics of the fluorescent species *Bt* and *Pf* was extracted from stacks of fluorescent images calculating the correlation coefficient, *r*_*c*_, between consecutive frames (Fig. 7A). *Bt* installation exhibited a correlation coefficient *r*_*c*_ <0.5 (Fig. 7A), attesting for a loose attachment on the surface corroborated by the colocalization maps of successive images (Fig. 7B). After about 2 hours, the correlation coefficient increased, which coincided with the formation of fluorescent asters resulting from the progressive aggregation of *Bt* around *Kv* cells which physically stabilized the connected *Bt* cells while the others remained highly mobile (Movie S3).

**Figure 7:**
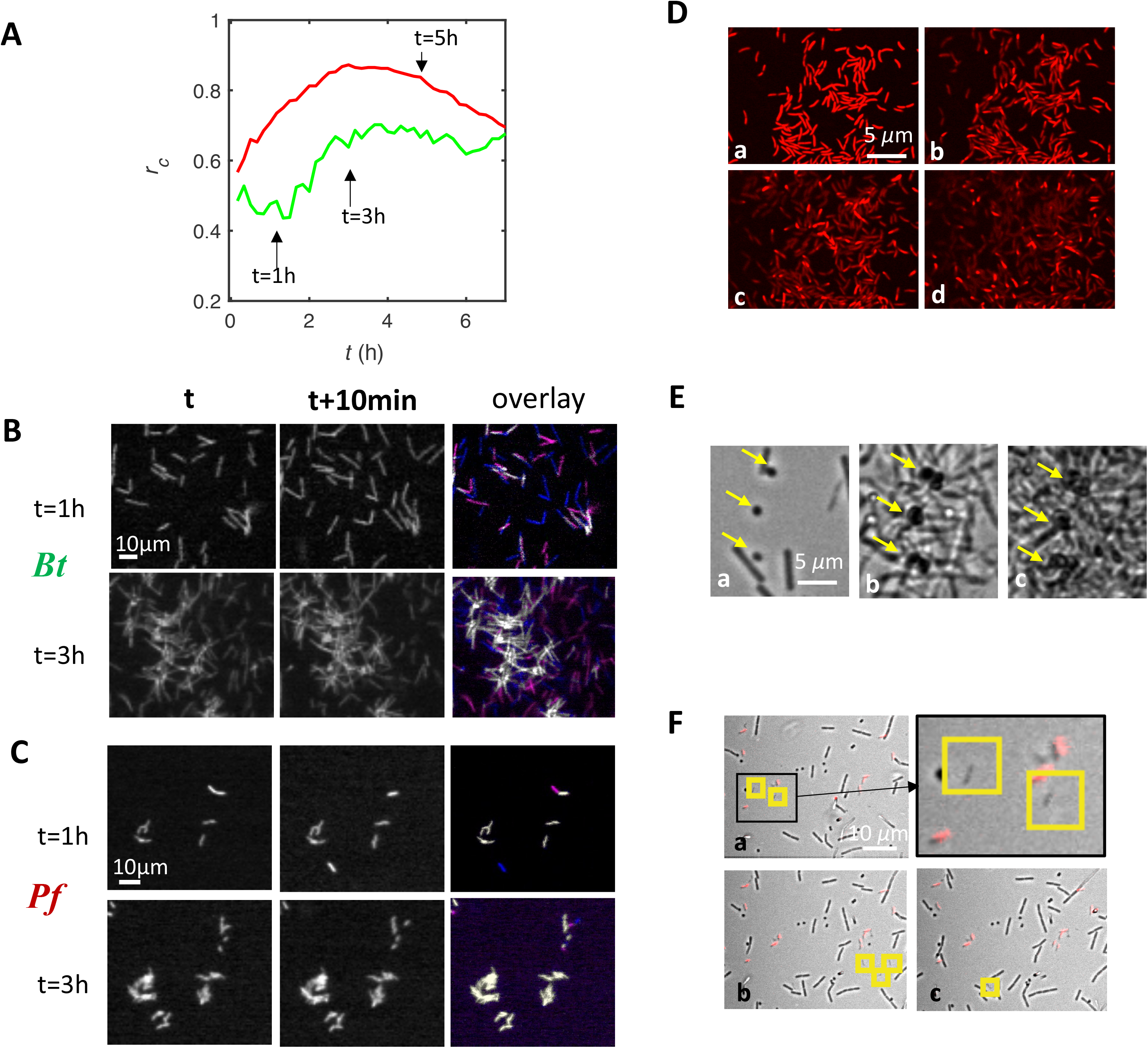
Species exhibit distinct local dynamics in the 4S community. **(A)** Consecutive frames correlation coefficient calculated from time-lapse recordings of *Bt*-FAST (green curve) and *Pf-*mCherry (red curve) in the 4S biofilm community. **(B, C)** Colocalization maps of *Bt* **(B)** and *Pf* **(C)** cells showing spatial self-overlap between time t and t’=t+10 min taken at t=1h (upper panels) and t=3h (lower panels). Light grey pixels correspond to non-moving cells (pixels unchanged between t and t’), blue pixels correspond to newly appeared cells (pixels dark at t and lighted at t’) and magenta pixels correspond to removed cells (pixels lighted at t and dark at t’). **(D)** Confocal images of *Pf*-mCherry in the 4S community taken at t= a) 6h40; b)7h; c)7h30 d)8h showing full detachment of the cell groups; see also colocalization maps in Fig. S6. **(E)** Transmitted light images of the 4S biofilm taken at time t = 1h (a) ; 12h (b) ; 37h (c) showing *Kv* (indicated by the yellow arrows) fixed localization all along the biofilm growth. **(F)** Successive frames (a,b,c) of the 4S biofilm in transmitted light and red fluorescence taken from t=1h (a) with a 10 mins interval. Upper right frame is a zoom in of (a). Yellow squares spot the *Rh* cells illustrating their high dynamics with different positions on each frame.

*Pf* cells exhibited a strong surface anchoring characterized by correlation coefficients comprised between 0.8 and 0.9 after 2 hours installation (Fig. 7A). The colocalization maps show that most of the initial attachments generated microcolonies (Fig. 7C). At time t≈5h, corresponding to the first climax of the development of *Pf* population on the bottom surface, *r*_*c*_ decreased while *Pf* cells simultaneously started to detach, shifting from firmly attached to essentially detached cells in less than two hours (Fig.7D and Fig. S6). Noticeably, a similar detachment occurred on the top surface but delayed by about 6 hours consistently with the shift of the climaxes mentioned above.

*Kv* dynamics was evaluated from the localization maps established to evaluate its spatial distribution. Stable attachment of isolated cells randomly dispersed over the whole bottom surface of the channel was observed from the first minutes of the biofilm initiation. As biofilm grew, compact clusters of a few cells formed essentially at the same location than the initial attachment (Fig. 7E). At least 90% of the *Kv* clusters came from single cell initial anchors. The correlation coefficient calculated from the position map was found equal to 0.94 (Fig. S7).

*Rh* was difficult to follow, exhibiting high dynamics. On the first images, a few cells could be visually detected. Their surface-dwelling time was found below the 10 min of the frame period, meaning *Rh* cells were never found at the same location on consecutive frames (Fig. 7F). Therefore, the correlation coefficient for *Rh* within the installation period was assigned to 0 as a principle.

The 4 species installation dynamics ranges from highly dynamical with *Rh* (*r*_*c*_=0) to completely fixed with *Kv* (*r*_*c*_ =0.95). The *r*_*c*_ values which measure the bacterial residence on the surface in the presence of the flow show the significantly distinct capacities of the 4 species to adhere on the surface. The correlation profile of *Bt* which r_*c*_ value increases upon its association with *Kv* also shows how physical interspecies interaction can improve the colonizing potential of poorly adhesive species.

### The 4-species combinatorics highlights community-specific traits

Next, we raised the question of which are the community-specific traits. To this purpose, we designed a combinatorial approach consisting in injecting all the possible sub-combinations of the 4 species in different parallel channels (14 in total). Then, we collected the optical signals of all the channels along the first 36 hours following channel seeding. We exhaustively show the mixes recordings in supplementary material (Fig. S8) but focus in the following on the features the most significant for the interpretation of the community traits.

#### Bt improves its installation and physical stability in the 4S

The biofilm built by *Bt* alone exhibited several characteristics distinct from the 4-species biofilm. In particular the initial population oscillation was absent, replaced by a 10 to 12 hours-long lag phase where the cell amount at the surface remained low (Fig. 8A panel a). We also observed that local dynamics were characterized by correlation coefficient values below 0.4 (Fig. 8B) which indicated that *Bt* cells essentially did not attach on the surface in the absence of the other species. After 10 hours, the fluorescence signal significantly increased reporting a colonization phase similar to the second phase displayed by *Bt* in the 4S biofilm but affected by a strong instability. The standard deviation was of the order of the signal itself due to massive random detachment events which resulted in important variations of the signal (Movie S4). In addition, the single-species *Bt*-FAST and *Bt*-GFP biofilms exhibited fully superimposable fluorescence signals which indicates no O_2_ depletion occurred in this situation (58)(Fig. S9). In the presence of *Pf, Bt* development profiles did not drastically changed compared to that of *Bt* alone, except the colonization phase appeared to be paused for a few hours compared to the expansion of *Bt* alone (Fig. 8A panel b). The standard deviation of the data sets was also 3 times less than for *Bt* alone, suggesting a physically more stable biofilm had formed. The biofilm built by the *Bt-Kv* pair (Fig. 8A panel c) confirmed the specific *Bt*-*Kv* physical interaction which led to the formation of *Bt* asters around *Kv* (Fig. 6A and Movie S3) and to the physical grip effect that promoted *Bt* early installation as showed by the increase of the fluorescence intensity and of the consecutive frames correlation coefficient (Fig. 8B). In the 3-species *Bt-Pf*-*Kv* community, the *Bt* development profile combined sequentially (i) the traits of the *Bt-Kv* pair: the initial promoted *Bt* growth, and (ii) the traits of the *Bt-Pf* pair: the second phase induced latency (Fig. 8A panel d). This 3-species community closely resembles the 4S community with the initial oscillation and a second growth phase. Yet, it should be mentioned that, in comparison with to *Bt-Pf* and 4S, the 3-species *Bt-Pf*-*Kv* community exhibited an extended *Bt* second phase latency.

**Figure 8:**
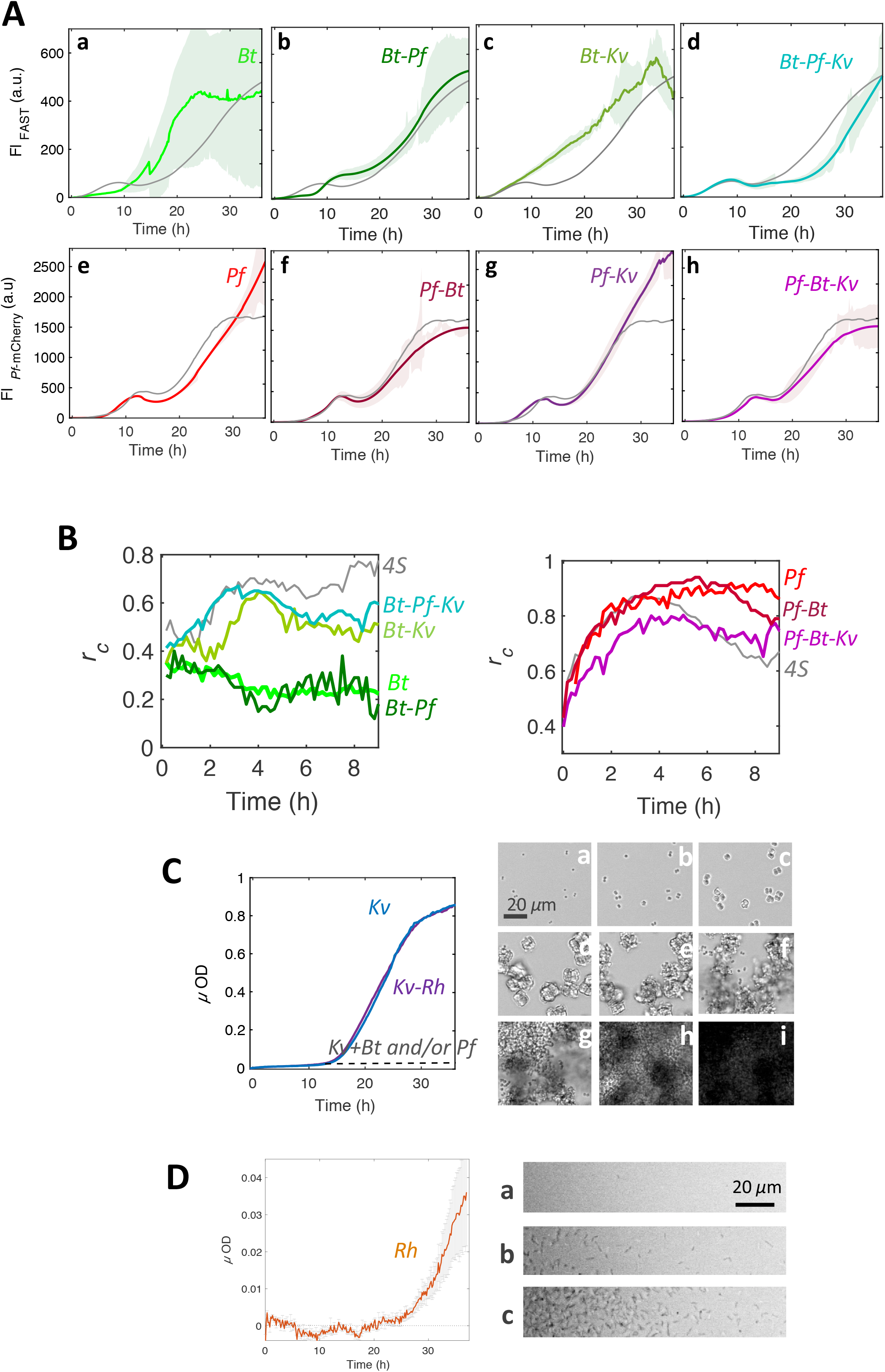
Species combinatorics highlights community specific-traits. (**A)** Fluorescence signals of *Bt* (a-d) and *Pf* (e-h) in various sub-combinations of the 4 species : single-species (a,e), pairs (b,c,f,g) and triplets (d,h). **(B)** Consecutive frames correlation coefficient for *Bt* (left panel) and *Pf* (right panel) for the different combinations mentioned on the figure. Experimental conditions other than the biofilm composition are as in Fig. 3. **(C)** Micro-optical density of *Kv* single-species biofilm (in blue) and *Kv-Rh* pair (in purple) (graph on the left); the dashed line shows the level of *Kv* development in the presence of *Bt*, alone, *Pf* alone and *Pf* and *Bt* together, deduced from *Kv* area determination and mean *µ*OD of a *Kv* cluster. Images on the right are snapshots of the single-species biofilm taken at time t = 10 min(a); 3h30(b); 9h(c); 15h(d); 16h30(e); 18h30(f); 23h30(g); 27h30(h); 30h(i). **(D)** Micro-optical density of *Rh* single-species biofilm (graph on the left) and corresponding snapshots taken at time t = 10h(a); 25h(b); 37h(c). The curve of the studied combination in each panel (color bold line) is the average of at least 3 independent samples and appears shaded with the standard deviation of the data set. The curve of the corresponding species in the 4S community is plotted in grey for the sake of comparison.

#### Pf undergoes a Bt-imposed dynamic equilibrium

*Pf* in the various combinations exhibited the same main traits evidenced in the 4S biofilm except the rise of the dynamic equilibrium which was absent in all the combinations devoid of *Bt* (Fig. 8A panel e-h). *Pf* appears as the less impacted species in the various combinations, simply forced to equilibrium by the presence of *Bt* after 30h of combined growth.

#### Kv development is inhibited in the 4S community

Injected alone in the channel, *Kv* population formed a profuse single-species biofilm. The cells immediately adhered on the bottom glass slide and multiplied *in situ* forming compact cubic clusters over the first 10 hours of the biofilm formation as described above in the 4S community (Fig. 8C). Then, whereas the *Kv* clusters were halted in this stage in the 4S community, they went on growing exponentially in the single-species mode, forming larger cohesive clusters of tens of cells before bursting again into single cells which were in part removed by the flow, and in part readdressed to the surface, efficiently forming a thick biofilm (Fig. 8C and Movie S5). We found that the *Kv* population was inhibited by the presence of *Bt* alone, *Pf* alone and both *Bt* and *Pf* as quantified on the images (Fig. 8C and Fig. S10) exactly the same way it happened in the 4S (Fig. S5). Noticeably, *Kv* cells extracted from two-species and three-species biofilms with *Bt* and/or *Pf* were recovered by plating on agar forming colonies, which indicated they were still viable.

#### Rh, the community neutral element

*Rh* alone exhibited poor colonization abilities. A thin layer of attached cells concentrated at the edge of the channel emerged after about 25 hours (Fig. 8D). Its contribution in mixed communities was always optically overwhelmed by that of the other species and could not be characterized in real time. Although their presence was attested by channel extraction counting, *Rh* cells did not alter any characteristics of the other species in multispecies communities (Fig. S8).

## Discussion

We report here a real-time analysis of a 4-species bacterial assembly, composed from a more complex natural community (50) and grown under constant flow of growth medium in a laboratory millifluidic device. The simple and defined structure of the setup provided a controlled environment, the heterogeneity of which was sufficient to allow behavioral complexity to emerge.

We show that the 4S assembly forms a surface-attached community displaying a dynamic equilibrium, with a biomass robust to physical perturbation after about thirty hours seeding. We find that the 4S community settles according to a deterministic mechanism involving timed spatiotemporal phases of expansion and recession. However, two of them, *Pf* and *Bt* clearly dominate the developmental program. Remarkably, while the four species coexisted on the bottom surface, a specialist niche emerged on the top surface, accommodating exclusively *Pf*, who escaped gravity and bottom surface assignation. Considering the channel dimensions (section of 1 mm^2^) and the flow rate (1 ml/h), characteristic times of 2 mins for advection and of 10 mins to a few hours for molecular diffusion in the height of the channel can be evaluated from the Stokes-Einstein equation (59). Thereby, bottom and top surfaces habitats can be regarded as independent as well as their respective biomes, reminding — although at a much smaller scale and limited range — the biogeographical sorting that occurs in nature (60).

On the bottom surface, *Pf* and *Bt* appeared closely coupled in time and space. Our results support two distinct mechanisms of coupling. The first one, governing the first 25 hours of the biofilm formation, mediated by the environment and interpreted as a coupled response to the depletion in O_2_ of the environment principally resulting from the *Pf* population growth. This hypothesis is supported by the comparison of *Bt*-GFP and *Bt*-FAST fluorescence signals in the 4S and the single-species communities. Consistently, *Pf* development kinetics exhibited very similar profiles in a 4S and in a single-species biofilm within the first 25 hours of the formation, suggesting a prevalent imprint of *Pf* on the environment. In this scenario, the O_2_ depletion of the environment is also assumed to drive *Pf* detachment and spatial transition to the channel edges. Indeed, we know from previous experiments that, due to PDMS permeability to O_2_, renewal of O_2_ supplies occurs principally at the channel edges in our device (51, 58), explaining the selective growth of the aerobes in the regions of higher O_2_ level where they compete for the same resource. Here, we highlight a mechanism in which the spatial distribution is governed by the shaping of the environment by one of the species. Such a resource-driven spatial organization might be a key principle of the emergence of spatial structure in complex biofilms. It has also been previously described in colonies of mixed genotypes of *Pseudomonas aeruginosa* (61). Yet, reciprocal influence of spatial distribution on social interactions has also been described (40), confirming no unequivocal link between social behavior and spatial structure can be established (21). Moreover, we bring evidence that the spatial structure evolves along the biofilm development. The expression of a *Bt-Pf* interaction is also found in the characteristics of the dynamic equilibrium established after thirty hours of 4S biofilm growth. From the combinatorial assembly of multispecies communities, it appears clearly that *Bt* forces *Pf* to a plateau resulting from balanced growth and detachment by contrast to the continuous expansion *Pf* exhibits in its pure biofilm. The presence of *Pf* also provides a better physical stability to the co-existing population of *Bt*, as shown by the significant decrease of the standard deviation for *Bt* signal in the presence of *Pf*, and the reduction of the number of spikes due to detachment events.

These changes occurring in the stability of the biofilm point to the question of the assembly of the extracellular matrix in the multispecies context which just starts to be investigated (62). Indeed, biofilm stability is closely related to extracellular matrix properties such as its degree of cross-linking (63). The larger spectrum of components brought together in multispecies environment might offer novel opportunities to form cross-links. In this hypothesis, the matrix assembly is a community activity which occurs extracellularly as shown in a previous study on *Candida albicans* biofilms (64). This provides an illustration of how spatial reorganization leads to the emergence of novel properties, reinforcing the view of the critical role of the spatial distribution on the biofilm properties (20, 22, 24, 65, 66).

*Bt* and *Kv* displayed a specific physical interaction enabling several *Bt* cells to attach to a *Kv* cluster and to take advantage from the strong affinity of *Kv* cells for the bottom surface of the channel, in particular in the initial phase of the colonization where it hardly forms transient ties with the surface. This interspecies physical attachment has been mainly overlooked in the analyses of multispecies community structures compared to social interactions mediated by diffusible factors such as quorum sensing or metabolite tradeoffs (46, 67–69) except in the special case of the oral biofilms (70). A better understanding of the impact of these specific bindings on the biological functions as well as additional knowledge about their diversity and their evolutionary profile might open new perspectives in the comprehension of multispecies adherent communities.

Besides, *Kv* population experiences a major limitation of its development in the mixed communities containing *Bt* alone, *Pf* alone and both *Bt* and *Pf*. As both *Bt* and *Pf* inhibit *Kv* biofilm development without killing it, it is tempting to postulate an environmental effect driven by the development of neighbors rather than a specific interspecies interaction. However, at this stage, we have no cues for the involved parameter, in particular the inhibition did not show any positional dependence ruling out a link to O_2_ supply as assumed to explain *Bt* and *Pf* behaviors.

*Rh* appears as the freewheeling partner of the community. This highly motile cell, proved to develop poor attachment when grown as a single-species biofilm, still multiplies and persists in the 4S community in an apparently fully neutral mode with no detectable interaction with the other species. It should be mentioned that we gained poor information about it since it could not be tracked in the multispecies communities. However, we decided to conserve it in the consortium as an example of coexistence without interactions, anticipating a possible role of this neutral element in disturbed environmental conditions.

We conclude that our composite biofilm reaches its dynamical equilibrium based on local cell concentration, environmental physico-chemical properties constantly reshaped by the community development itself, and a deep correction of the species fitness by surface attachment. From a conceptual perspective, this community could be considered as individualistic to the sense of Gleason (71) but also as a limiting case of an organismic continuum as theoretically proposed by Liautaud et al. (72).

This multispecies biofilm emerges as the result of chance (a given species in a given environment) and necessity (individual species adaptation) deterministically leading to a unique community. The emergent properties could not be predicted from individual traits, involving a form of biofilm sociobiology which does not require that cells communicate with one another using specialized signaling molecules (73).

The operational experimental model and assembly principles we present here should prove useful to investigate multispecies bacterial community response to perturbations or elaborate new strategies (65) to manipulate biofilm functions in several fields including health, agriculture and environment.

## Acknowledgments

Authors would like to thank Pierre Nicolas and Cyprien Guérin for fruitful discussions and Carounagarane Doré for technical assistance in the devices elaboration.

The work was supported by a grant from French Agence Nationale pour la Recherche (ANR-15-CE02-0001-01 ACToP) and a MESRI fellowship to WBY.

## Notes

**Competing Interests**The authors declare there are no competing interests

### Competing Interest Statement

The authors have declared no competing interest.

## References

1. Falkowski PG, Fenchel T, Delong EF. The microbial engines that drive Earth’s biogeochemical cycles. Science. 2008;320(5879):1034–9.

2. Battin TJ, Besemer K, Bengtsson MM, Romani AM, Packmann AI. The ecology and biogeochemistry of stream biofilms. Nat Rev Microbiol. 2016;14(4):251–63.

3. Benton TG, Solan M, Travis JM, Sait SM. Microcosm experiments can inform global ecological problems. Trends Ecol Evol. 2007;22(10):516–21.

4. Faust K, Raes J. Microbial interactions: from networks to models. Nat Rev Microbiol. 2012;10(8):538–50.

5. Ponomarova O, Patil KR. Metabolic interactions in microbial communities: untangling the Gordian knot. Curr Opin Microbiol. 2015;27:37–44.

6. Strom SL. Microbial ecology of ocean biogeochemistry: a community perspective. Science. 2008;320(5879):1043–5.

7. Morris BEL, Henneberger R, Huber H, Moissl-Eichinger C. Microbial syntrophy: interaction for the common good. FEMS Microbiology Reviews. 2013;37(3):384–406.

8. Moscoviz R, Flayac C, Desmond-Le Quemener E, Trably E, Bernet N. Revealing extracellular electron transfer mediated parasitism: energetic considerations. Sci Rep. 2017;7(1):7766.

9. Watkins ER, Maiden MC, Gupta S. Metabolic competition as a driver of bacterial population structure. (1746-0921 (Electronic)).

10. Hart SP, Usinowicz J, Levine JM. The spatial scales of species coexistence. Nat Ecol Evol. 2017;1(8):1066–73.

11. Jessup CM, Kassen R, Forde SE, Kerr B, Buckling A, Rainey PB, et al. Big questions, small worlds: microbial model systems in ecology. Trends Ecol Evol. 2004;19(4):189–97.

12. Konopka A, Lindemann S, Fredrickson J. Dynamics in microbial communities: unraveling mechanisms to identify principles. ISME J. 2015;9(7):1488–95.

13. Gause GF. The struggle for existence. Baltimore: The Williams & Wilkins company; 1934 Dec, 1934.

14. Geesey GG. Bacterial behavior at surfaces. Curr Opin Microbiol. 2001;4(3):296–300.

15. Flemming HC, Wingender J, Szewzyk U, Steinberg P, Rice SA, Kjelleberg S. Biofilms: an emergent form of bacterial life. Nat Rev Microbiol. 2016;14(9):563–75.

16. Hall-Stoodley L, Costerton JW, Stoodley P. Bacterial biofilms: from the natural environment to infectious diseases. Nat Rev Microbiol. 2004;2(2):95–108.

17. Costerton JW. Overview of microbial biofilms. J Ind Microbiol. 1995;15(3):137–40.

18. van Gestel J, Kolter R. When We Stop Thinking about Microbes as Cells. J Mol Biol. 2019;431(14):2487–92.

19. Stewart PS, Franklin MJ. Physiological heterogeneity in biofilms. Nat Rev Microbiol. 2008;6(3):199–210.

20. Kim HJ, Boedicker JQ, Choi JW, Ismagilov RF. Defined spatial structure stabilizes a synthetic multispecies bacterial community. Proc Natl Acad Sci U S A. 2008;105(47):18188–93.

21. Nadell CD, Drescher K, Foster KR. Spatial structure, cooperation and competition in biofilms. Nat Rev Microbiol. 2016;14(9):589–600.

22. Bridier A, Piard JC, Pandin C, Labarthe S, Dubois-Brissonnet F, Briandet R. Spatial Organization Plasticity as an Adaptive Driver of Surface Microbial Communities. Front Microbiol. 2017;8:1364.

23. Cutler NA, Chaput DL, Oliver AE, Viles HA. The spatial organization and microbial community structure of an epilithic biofilm. FEMS Microbiol Ecol. 2015;91(3).

24. France MT, Forney LJ. The Relationship between Spatial Structure and the Maintenance of Diversity in Microbial Populations. Am Nat. 2019;193(4):503–13.

25. Azeredo J, Azevedo NF, Briandet R, Cerca N, Coenye T, Costa AR, et al. Critical review on biofilm methods. Crit Rev Microbiol. 2017;43(3):313–51.

26. Neu TR, Manz B, Volke F, Dynes JJ, Hitchcock AP, Lawrence JR. Advanced imaging techniques for assessment of structure, composition and function in biofilm systems. FEMS Microbiol Ecol. 2010;72(1):1–21.

27. Wessel AK, Hmelo L, Parsek MR, Whiteley M. Going local: technologies for exploring bacterial microenvironments. Nat Rev Microbiol. 2013;11(5):337–48.

28. Rusconi R, Garren M, Stocker R. Microfluidics expanding the frontiers of microbial ecology. Annu Rev Biophys. 2014;43:65–91.

29. Burmeister A, Hilgers F Fau - Langner A, Langner A Fau - Westerwalbesloh C, Westerwalbesloh C Fau - Kerkhoff Y, Kerkhoff Y Fau - Tenhaef N, Tenhaef N Fau - Drepper T, et al. A microfluidic co-cultivation platform to investigate microbial interactions at defined microenvironments. (1473-0189 (Electronic)).

30. Moons P, Michiels CW, Aertsen A. Bacterial interactions in biofilms. Crit Rev Microbiol. 2009;35(3):157–68.

31. Niu B, Kolter R. Quantification of the Composition Dynamics of a Maize Root-associated Simplified Bacterial Community and Evaluation of Its Biological Control Effect. Bio Protoc. 2018;8(12).

32. Alnahhas RN, Winkle JJ, Hirning AJ, Karamched B, Ott W, Josic K, et al. Spatiotemporal Dynamics of Synthetic Microbial Consortia in Microfluidic Devices. ACS Synth Biol. 2019;8(9):2051–8.

33. Liu W, Roder HL, Madsen JS, Bjarnsholt T, Sorensen SJ, Burmolle M. Interspecific Bacterial Interactions are Reflected in Multispecies Biofilm Spatial Organization. Front Microbiol. 2016;7:1366.

34. Momeni B, Brileya KA, Fields MW, Shou W. Strong inter-population cooperation leads to partner intermixing in microbial communities. Elife. 2013;2:e00230.

35. Ratzke C, Gore J. Self-organized patchiness facilitates survival in a cooperatively growing Bacillus subtilis population. Nat Microbiol. 2016;1:16022.

36. Ren D, Madsen JS, Sorensen SJ, Burmolle M. High prevalence of biofilm synergy among bacterial soil isolates in cocultures indicates bacterial interspecific cooperation. ISME J. 2015;9(1):81–9.

37. Almeida C, Azevedo NF, Santos S, Keevil CW, Vieira MJ. Discriminating multi-species populations in biofilms with peptide nucleic acid fluorescence in situ hybridization (PNA FISH). PLoS One. 2011;6(3):e14786.

38. Benoit MR, Conant CG, Ionescu-Zanetti C, Schwartz M, Matin A. New device for high-throughput viability screening of flow biofilms. Appl Environ Microbiol. 2010;76(13):4136–42.

39. Roder HL, Liu W, Sorensen SJ, Madsen JS, Burmolle M. Interspecies interactions reduce selection for a biofilm-optimized variant in a four-species biofilm model. Environ Microbiol Rep. 2019;11(6):835–9.

40. Liu W, Russel J, Burmolle M, Sorensen SJ, Madsen JS. Micro-scale intermixing: a requisite for stable and synergistic co-establishment in a four-species biofilm. ISME J. 2018;12(8):1940–51.

41. Malic S, Hill KE, Hayes A, Percival SL, Thomas DW, Williams DW. Detection and identification of specific bacteria in wound biofilms using peptide nucleic acid fluorescent in situ hybridization (PNA FISH). Microbiology. 2009;155(Pt 8):2603–11.

42. Valm AM, Mark Welch JL, Rieken CW, Hasegawa Y, Sogin ML, Oldenbourg R, et al. Systems-level analysis of microbial community organization through combinatorial labeling and spectral imaging. Proc Natl Acad Sci U S A. 2011;108(10):4152–7.

43. Costa AM, Mergulhao FJ, Briandet R, Azevedo NF. It is all about location: how to pinpoint microorganisms and their functions in multispecies biofilms. Future Microbiol. 2017;12:987–99.

44. Wagner M, Nielsen PH, Loy A, Nielsen JL, Daims H. Linking microbial community structure with function: fluorescence in situ hybridization-microautoradiography and isotope arrays. Curr Opin Biotechnol. 2006;17(1):83–91.

45. Mattei MR, Frunzo L, D’Acunto B, Pechaud Y, Pirozzi F, Esposito G. Continuum and discrete approach in modeling biofilm development and structure: a review. J Math Biol. 2018;76(4):945–1003.

46. Borenstein DB, Meir Y, Shaevitz JW, Wingreen NS. Non-local interaction via diffusible resource prevents coexistence of cooperators and cheaters in a lattice model. PLoS One. 2013;8(5):e63304.

47. Bridier A, Briandet R, Bouchez T, Jabot F. A model-based approach to detect interspecific interactions during biofilm development. Biofouling. 2014;30(7):761–71.

48. Xavier JB, Martinez-Garcia E, Foster KR. Social evolution of spatial patterns in bacterial biofilms: when conflict drives disorder. Am Nat. 2009;174(1):1–12.

49. Kreft JU, Picioreanu C, Wimpenny JW, van Loosdrecht MC. Individual-based modelling of biofilms. Microbiology. 2001;147(Pt 11):2897–912.

50. Mettler E, Carpentier, B.. Location, enumeration and identification of the microbial contamination after cleaning of EPDM gaskets introduced into a milk pasteurization line. Dairy Science and Technology. 1997;77:489–503.

51. Thomen P, Robert J, Monmeyran A, Bitbol AF, Douarche C, Henry N. Bacterial biofilm under flow: First a physical struggle to stay, then a matter of breathing. PLoS ONE. 2017;12(4):e0175197.

52. Sheppard AE, Poehlein A, Rosenstiel P, Liesegang H, Schulenburg H. Complete Genome Sequence of Bacillus thuringiensis Strain 407 Cry. Genome Announc. 2013;1(1):e00158–12.

53. Plamont MA, Billon-Denis E, Maurin S, Gauron C, Pimenta FM, Specht CG, et al. Small fluorescence-activating and absorption-shifting tag for tunable protein imaging in vivo. Proc Natl Acad Sci U S A. 2016;113(3):497–502.

54. Lagendijk EL, Validov S, Lamers GE, de Weert S, Bloemberg GV. Genetic tools for tagging Gram-negative bacteria with mCherry for visualization in vitro and in natural habitats, biofilm and pathogenicity studies. FEMS Microbiol Lett. 2010;305(1):81–90.

55. Schneider CA, Rasband WS, Eliceiri KW. NIH Image to ImageJ: 25 years of image analysis. Nat Methods. 2012;9(7):671–5.

56. Gonze D, Coyte KZ, Lahti L, Faust K. Microbial communities as dynamical systems. Curr Opin Microbiol. 2018;44:41–9.

57. Coyte KZ, Schluter J, Foster KR. The ecology of the microbiome: Networks, competition, and stability. Science. 2015;350(6261):663–6.

58. Monmeyran A, Thomen P, Jonquiere H, Sureau F, Li C, Plamont MA, et al. The inducible chemical-genetic fluorescent marker FAST outperforms classical fluorescent proteins in the quantitative reporting of bacterial biofilm dynamics. Sci Rep. 2018;8(1):10336.

59. Hynes JT. Statistical Mechanics of Molecular Motion in Dense Fluids. Annual Review of Physical Chemistry. 1977;28(1):301–21.

60. Monard C, Gantner S, Bertilsson S, Hallin S, Stenlid J. Habitat generalists and specialists in microbial communities across a terrestrial-freshwater gradient. Sci Rep. 2016;6:37719.

61. Mitri S, Clarke E, Foster KR. Resource limitation drives spatial organization in microbial groups. ISME J. 2016;10(6):1471–82.

62. Karygianni L, Ren Z, Koo H, Thurnheer T. Biofilm Matrixome: Extracellular Components in Structured Microbial Communities. Trends Microbiol. 2020;28(8):668–81.

63. Galy O, Latour-Lambert P, Zrelli K, Beloin C, Ghigo JM, Henry N. Mapping of bacterial biofilm local mechanics by magnetic microparticle actuation. Biophysical journal. 2012;5(6):1400–8.

64. Mitchell KF, Zarnowski R, Sanchez H, Edward JA, Reinicke EL, Nett JE, et al. Community participation in biofilm matrix assembly and function. Proc Natl Acad Sci U S A. 2015;112(13):4092–7.

65. Ben Said S, Tecon R, Borer B, Or D. The engineering of spatially linked microbial consortia - potential and perspectives. Curr Opin Biotechnol. 2020;62:137–45.

66. Nadell CD, Foster KR, Xavier JB. Emergence of spatial structure in cell groups and the evolution of cooperation. PLoS Comput Biol. 2010;6(3):e1000716.

67. Mukherjee S, Bassler BL. Bacterial quorum sensing in complex and dynamically changing environments. Nat Rev Microbiol. 2019;17(6):371–82.

68. An D, Danhorn T, Fuqua C, Parsek MR. Quorum sensing and motility mediate interactions between Pseudomonas aeruginosa and Agrobacterium tumefaciens in biofilm cocultures. Proc Natl Acad Sci U S A. 2006;103(10):3828–33.

69. Moons P, Van Houdt R, Aertsen A, Vanoirbeek K, Michiels CW. Quorum sensing dependent production of antimicrobial component influences establishment of E. coli in dual species biofilms with Serratia plymuthica. Commun Agric Appl Biol Sci. 2005;70(2):195–8.

70. Bowen WH, Burne RA, Wu H, Koo H. Oral Biofilms: Pathogens, Matrix, and Polymicrobial Interactions in Microenvironments. Trends Microbiol. 2018;26(3):229–42.

71. McIntosh RP. H. A. Gleason’s ‘individualistic concept’ and theory of animal communities: a continuing controversy. Biol Rev Camb Philos Soc. 1995;70(2):317–57.

72. Liautaud K, van Nes EH, Barbier M, Scheffer M, Loreau M. Superorganisms or loose collections of species? A unifying theory of community patterns along environmental gradients. Ecol Lett. 2019;22(8):1243–52.

73. Nadell CD, Xavier JB, Foster KR. The sociobiology of biofilms. FEMS Microbiol Rev. 2009;33(1):206–24.

